# Single-molecule live-cell RNA imaging with CRISPR-Csm

**DOI:** 10.1101/2024.07.14.603457

**Authors:** Chenglong Xia, David Colognori, Xueyang Jiang, Ke Xu, Jennifer A. Doudna

## Abstract

High-resolution, real-time imaging of RNA is essential for understanding the diverse, dynamic behaviors of individual RNA molecules in single cells. However, single-molecule live-cell imaging of unmodified endogenous RNA has not yet been achieved. Here, we present single-molecule live-cell fluorescence *in situ* hybridization (smLiveFISH), a robust approach that combines the programmable RNA-guided, RNA-targeting CRISPR-Csm complex with multiplexed guide RNAs for efficient, direct visualization of single RNA molecules in a range of cell types, including primary cells. Using smLiveFISH, we tracked individual endogenous *NOTCH2* and *MAP1B* mRNA transcripts in living cells and identified two distinct localization mechanisms: co-translational translocation of *NOTCH2* mRNA at the endoplasmic reticulum, and directional transport of *MAP1B* mRNA toward the cell periphery. This method has the potential to unlock principles governing the spatiotemporal organization of native transcripts in health and disease.

## Introduction

RNA is directly involved in protein synthesis and regulates gene expression at both transcriptional and post-transcriptional levels^1^. Beyond RNA sequence, the spatial and temporal dynamics of individual transcripts control these activities. Dynamic and orchestrated interactions between RNA, RNA-binding proteins (RBPs) and other cellular machinery occur at particular subcellular regions and timepoints^2–4^. For example, zipcode-binding protein 1 (ZBP1) mediates directional transport of β-actin mRNA from the nucleus to the leading edge of fibroblasts^5^, and elongation factor 1α (EF1α) subsequently anchors these transcripts to actin filaments at the cell edge for local translation^6^.

Live-cell RNA imaging methods have begun to reveal RNA dynamics within individual cells, highlighting the value of such interrogations^2–4^. However, stem-loop^7, 8^ or aptamer^9^ labeling methods require the genetic insertion of sequences within specific regions of RNA or rely on exogenous expression of tagged RNA^7–9^, manipulations that are both time-consuming and can interfere with native RNA behavior^10^. Approaches to visualize unmodified endogenous RNA based on molecular beacons^11^ or clustered regularly interspaced short palindromic repeats (CRISPR)-CRISPR-associated (Cas) proteins (CRISPR–Cas) systems^12–15^ have limited single-molecule resolution (often restricted to highly abundant, repetitive RNAs)^3, 4^, and molecular beacons additionally suffer from excessive background signals caused by endosome-entrapped probes^3, 4^.

Here we describe single-molecule live-cell fluorescence *in situ* hybridization (smLiveFISH), an alternative strategy for visualizing any unmodified endogenous transcript. Using the RNA-targeting type III-A CRISPR–Csm system from *S. thermophilus* with multiplexed guide RNAs, smLiveFISH can track individual mRNA molecules in different types of living cells. We used smLiveFISH to analyze the behavior of individual endogenous mRNAs, *NOTCH2* and *MAP1B*, that encode a cell-surface receptor protein and a microtubule-associated protein, respectively. We found that *NOTCH2* mRNAs comprise two populations with distinct dynamics associated with co-translational polypeptide translocation across the endoplasmic reticulum (ER). In contrast, *MAP1B* mRNAs exhibit several distinct behaviors including directional transport in a translation-independent manner. We further show that smLiveFISH can detect differences in single-transcript localization in response to small molecules, underscoring the utility of assessing individual endogenous RNA behavior.

## Results

### Design and characterization of smLiveFISH

CRISPR-Cas complexes are programmable DNA or RNA nucleases from prokaryotic adaptive defense systems against bacteriophages^16, 17^. Cas nucleases bind to CRISPR RNAs (crRNAs) to form ribonucleoprotein (RNP) complexes that recognize nucleic acid targets using base-pairing complementarity between the crRNA and target DNA or RNA^16, 17^. Fluorescently labeled, catalytically inactivated Cas nucleases from different types of CRISPR-Cas systems (such as RCas9^12^, dCas13^13, 14^, and dCsm^15^) have been used to label RNAs of interest in live cells. However, these approaches have yet to achieve single-molecule resolution because each RNA is targeted with a single crRNA, resulting in a similar fluorescence intensity between free and target-bound RNPs. Only RNA granules or RNAs with repeated sequences produce sufficient signal from multiple copies of bound RNPs that is distinguishable from non-specific binding or background signal.

To overcome this signal-to-noise issue, we drew inspiration from single-molecule fluorescence *in situ* hybridization^18, 19^ (smFISH). In smFISH^18, 19^, short, fluorescently labeled DNA probes are tiled along the target mRNA, increasing the signal-to-noise ratio and allowing detection by microscopy of equal-intensity diffraction-limited spots in fixed cells^18, 19^. We reasoned that single-molecule imaging of endogenous RNAs in living cells could be possible if fluorescently tagged RNPs can be simultaneously tiled along a target RNA (Fig. 1a).

**Fig. 1:**
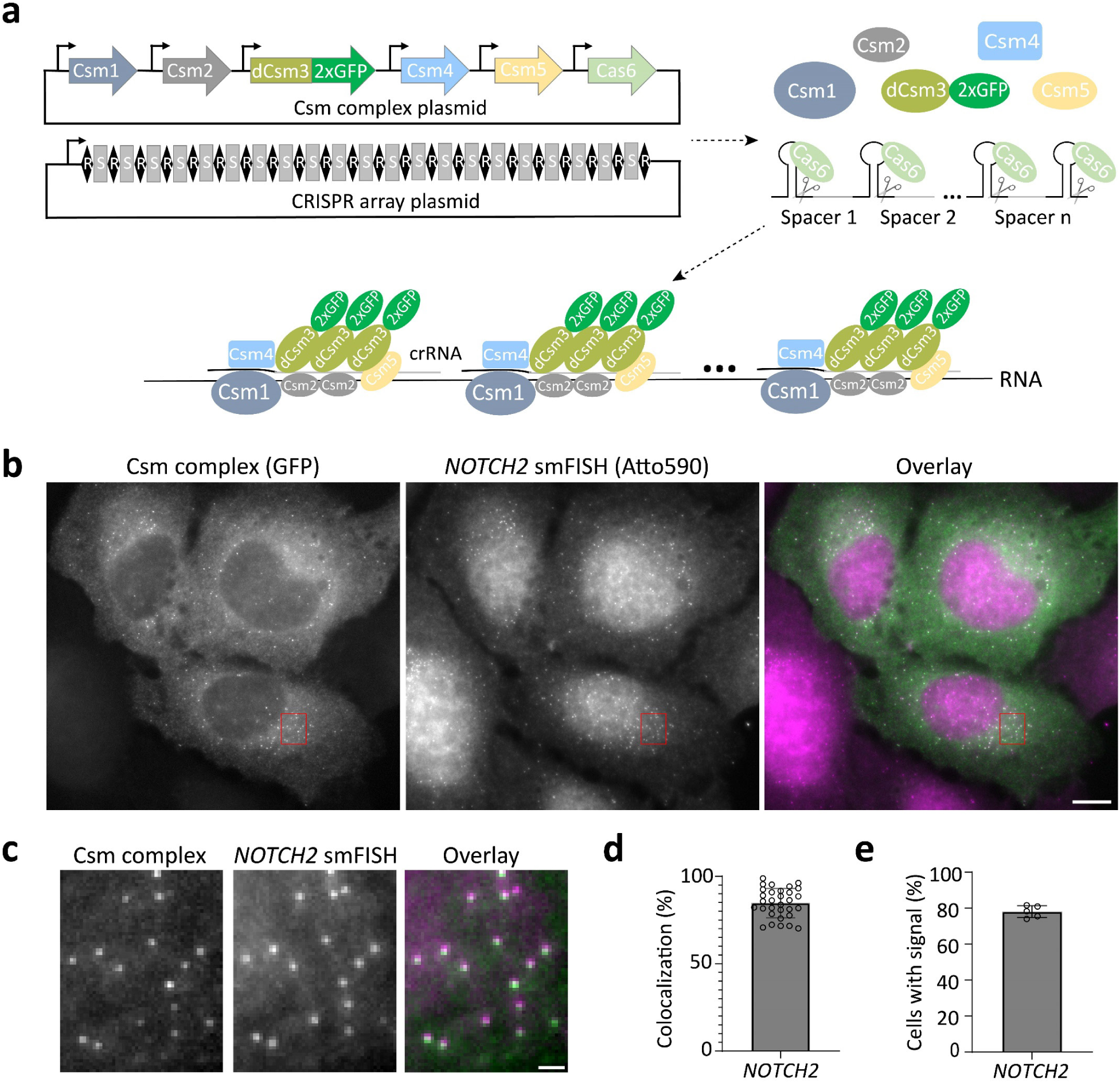
Imaging native single mRNA molecules with smLiveFISH. **a**, Schematic of the smLiveFISH system using multiplexed guides against a target RNA to achieve single-molecule resolution. Upon transfection with Csm and CRISPR array plasmids, cells produce Csm1, Csm2, dCsm3-2xGFP, Csm4, Csm5, and Cas6 proteins along with the pre-crRNA array. Cas6 processes the pre-crRNA array into individual crRNAs that assemble with Csm proteins into RNPs. RNPs, each with their own crRNA spacer, bind target RNA molecules simultaneously via base-pair complementarity, allowing RNA detection at single-molecule resolution. **b**, Fixed-cell image of individual *NOTCH2* mRNAs labeled by GFP-tagged Csm complex and 48 *NOTCH2*-targeting spacers (left), image of individual *NOTCH2* mRNAs labeled by smFISH probes (middle), and their overlay (right). Scale bar, 10 μm. **c**, Enlarged view of the red boxed region in **b.** Scale bar, 1 μm. **d**, Percent colocalization of Csm complex foci and smFISH foci (measured as Csm complex foci colocalized with smFISH foci divided by Csm complex foci per cell). Error bar indicates mean ± s.d., each dot represents one cell, n=31 cells. **e**, Percentage of transfected cells with Csm-complex-labeled foci. Images obtained from five randomly selected 7x7 tiling regions from three biological replicates (n=275 cells). Error bar indicates mean ± s.d., each dot represents one 7x7 tiling region.

For initial studies we compared GFP-fused Cas13 and Csm complex, both of which can generate individual crRNAs by processing pre-crRNAs, as catalyzed by Cas13^20, 21^ and Cas6^22^, respectively. To evaluate the labeling performance of dPspCas13b^14^, dRfxCas13d^21^ and dCsm proteins^15^, we targeted *XIST* RNA, a long noncoding transcript that forms large clouds from hundreds of *XIST* copies in HEK293T cell nuclei^23^. We used a single crRNA with eight complementary target sites on a repetitive region of *XIST*^15^ (Supplementary Table 1), and validated labeling with H2AK119ub, a heterochromatin mark enriched at the inactivated X chromosome which overlaps with *XIST* RNA. Only the Csm complex could label *XIST* robustly, an observation consistent with prior results^15, 24^ (Extended Data Fig. 1). Notably, the Csm system has other advantages relative to Cas13. First, it contains multiple GFP-linked catalytically inactive Csm3 molecules^15, 25^ (≥3 per complex) (Fig. 1a), which enhances signal and may aid in single-molecule detection. Second, it has higher binding affinity for RNA (K_d_ = 0.3 nM ^22^) relative to that for Cas13 (K_d_ = ∼10 nM^26^). Based on these observations, we focused on the Csm complex to develop a single-molecule mRNA labeling system.

To target mRNAs in the cytoplasm, we removed the nuclear localization signal (NLS) from each protein component of the Csm complex^15^ and encoded them within a single plasmid (Fig. 1a, Supplementary Table 2). To facilitate the export of pre-crRNA from the nucleus to the cytoplasm, we added a short signal sequence^27^ present in naturally intron-less mRNAs to the 5’ end of the pre-crRNA array and expressed it from a CAG (Pol II) promoter (Supplementary Table 2). We selected *NOTCH2* mRNA as a cytoplasmic target for proof of concept for two reasons. First, its length permits the design of distinct smFISH probes to validate labeling by the Csm complex. Second, it encodes a cell membrane protein and is thus enriched near the endoplasmic reticulum (ER)^28–30^, which is useful for RNA-centric exploration of co-translational protein translocation. In each of two crRNA array plasmids developed for this experiment (Extended Data Fig. 2, Supplementary Table 2), all 24 36-nt targeting sequences were designed to bind in a tiled fashion along the length of the 3’UTR of *NOTCH2* mRNA to avoid interfering with mRNA translation^31^.

While crRNA arrays have been used for multiplexed targeting in human cells, their length was relatively short (generally <10 crRNAs), and their processing was not directly demonstrated^15, 32^. To test whether the Csm-associated Cas6 endonuclease could process long crRNA arrays in cells (Fig. 1a), we used a single FISH probe (Extended Data Fig. 3a, Supplementary Table 3) complementary to the direct repeats to detect individual pre-crRNA arrays in U2OS cells. We observed high expression of pre-crRNA array transcripts, indicated by diffraction-limited puncta, in both the cytoplasm and nucleus of U2OS cells co-transfected with a crRNA array-encoding plasmid and an empty vector expressing GFP alone (Extended Data Fig. 3a). Upon co-transfection of plasmids encoding the crRNA array and the GFP-tagged Csm complex the individual spots disappeared from the cytoplasm (where Cas6 and Csm proteins are localized), consistent with pre-crRNA array processing (Extended Data Fig. 3b). Additionally, in cells co-transfected with plasmids encoding the crRNA array and Csm-GFP, single-molecule spots were observed in the cytoplasm using GFP fluorescence detection, representing putative *NOTCH2* mRNA signals (Extended Data Fig. 3b).

### SmLiveFISH enables visualization of individual endogenous *NOTCH2* mRNAs

We next utilized *NOTCH2* smFISH to identify the spots observed in cells transfected with crRNA array-encoding and Csm complex-encoding plasmids. Two-color imaging revealed strong colocalization of Csm labeled foci with smFISH spots, indicating that GFP-tagged Csm complexes successfully labeled endogenous *NOTCH2* mRNA (Fig. 1b-c). Quantification showed that 85% of Csm-labeled spots colocalized with smFISH spots (Fig. 1d). In addition, we found that 78% of transfected cells had clearly distinguishable spots in the cytoplasm, consistent with a high labeling efficiency (Fig. 1e). We next applied this method in other cell lines, including HEK293T, HeLa, primary human fibroblast IMR-90 and African green monkey COS-7—the latter having 94% *NOTCH2* 3’ UTR sequence identity to the human sequence. We observed robust labeling of endogenous *NOTCH2* mRNA in all of these cell types (Extended Data Fig. 4a-d). As controls, expression of the Csm complex alone only generated homogenous signal when GFP fluorescence was monitored (Extended Data Fig. 5a), and expression of the crRNA array alone exhibited only weak homogenous GFP autofluorescence (Extended Data Fig. 5b). In summary, these results demonstrate smLiveFISH to be a robust and efficient RNA imaging tool for visualizing unmodified endogenous RNA at single-molecule resolution across different cell types.

### *NOTCH2* mRNAs display translation-dependent population dynamics

Next, we conducted more comprehensive live-cell imaging of *NOTCH2* mRNA in U2OS cells and observed that *NOTCH2* mRNAs were divided into two distinct populations based on their diffusion dynamics: slow movement and fast movement (Fig. 2a-b). We reasoned that anchoring during co-translational translocation of *NOTCH2* nascent polypeptide across the ER membrane^28, 30^ could explain why one population of *NOTCH2* mRNA is nearly static. To test this possibility, we treated cells with puromycin, a translation elongation inhibitor that causes release of mRNA from the nascent polypeptide^33^. Upon treatment, the static population of *NOTCH2* mRNA rapidly decreased (Fig. 2c-d, Supplementary Movie 1). By single-molecule displacement/diffusivity mapping (SMdM) analysis^34^ of the two populations using a two-component diffusion mode (eqn. 2 in Methods), we found that puromycin treatment correlated with a shift in the *NOTCH2* mRNA population from slow to fast movement (Fig. 2e-f). Taken together, these results suggest that *NOTCH2* mRNA stationary binding to the perinuclear region is translation-dependent, consistent with mRNA docking for peptide translocation across the ER (Fig. 2g).

**Fig. 2:**
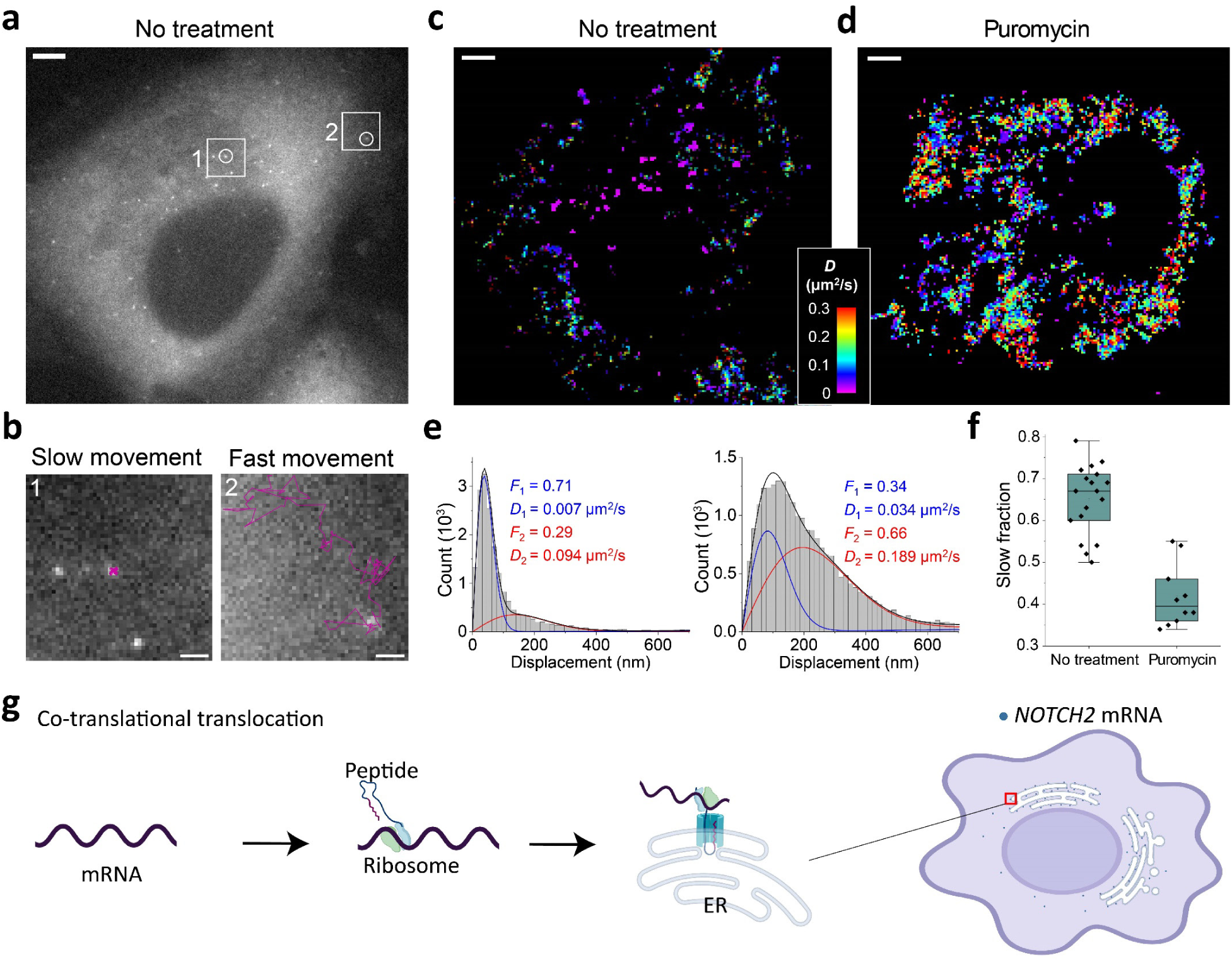
Dynamics of individual *NOTCH2* mRNAs in live cells. **a**, Single snapshot of individual *NOTCH2* mRNAs in live U2OS cell, scale bar, 5 μm. **b**, Movement trajectories (magenta) over time of two highlighted mRNA foci in **a**, reflecting slow and fast movements. Images were recorded at 10 frames per second over 25 seconds. Scale bar, 1 μm. **c**, Color-coded diffusivity map based on single-molecule displacement/diffusivity mapping (SMdM) analysis of *NOTCH2* mRNA foci in the cell shown in **a**. **d**, Same as **c**, but the cell was treated with puromycin for 1 h. Full movie shown in Supplementary Movie 1. **e**, Distributions of single-molecule displacements across successive frames for the data shown in **c,d** (histograms) and fits (curves) to a two-component diffusion mode (eqn 2). Blue curve: slow component; red curve: fast component; black curve: sum. Resultant fractions of the two components and *D* values are marked in the plots. **f**, Statistics of the slow fraction for treated and untreated cells. Each data point corresponds to the analysis result from one cell. **g**, A proposed co-translational translocation mechanism directing *NOTCH2* mRNA to the cell perinuclear region based on anchoring to the ER.

### *MAP1B* mRNA localization to the cell edge relies on directional transport

To explore the programmability of smLiveFISH as an RNA imaging platform, we examined *MAP1B* mRNA, which encodes a microtubule-associated protein involved in axon growth during neuronal development^35^. We hypothesized that *MAP1B* transcripts might employ a distinct mechanism of transport due to the different spatial localization pattern observed in fixed cells^29^ relative to ER-associated *NOTCH2* mRNA.

We designed 48 crRNAs tiling the 3’UTR of *MAP1B* mRNA (Supplementary Table 2) and transfected U2OS cells with Csm complex-encoding and crRNA array-encoding plasmids. Similar to *NOTCH2* mRNA labeling, single-molecule spots were observed in the cytoplasm using GFP fluorescence detection, representing putative *MAP1B* mRNA signals (Fig. 3a-b). We validated these spots with separate *MAP1B* smFISH probes bearing a second color and observed strong colocalization of Csm-labeled spots with smFISH spots (Fig. 3a-b). Using smLiveFISH, we investigated the spatial distribution of the labeled RNA species. By measuring the distance from mRNA molecules to the cell nucleus and/or cell edge, we found that *MAP1B* mRNAs are enriched at the cell periphery while *NOTCH2* mRNAs are enriched at the perinuclear region (Fig. 3c-d), in agreement with previous smFISH results in fixed cells^29^.

**Fig. 3:**
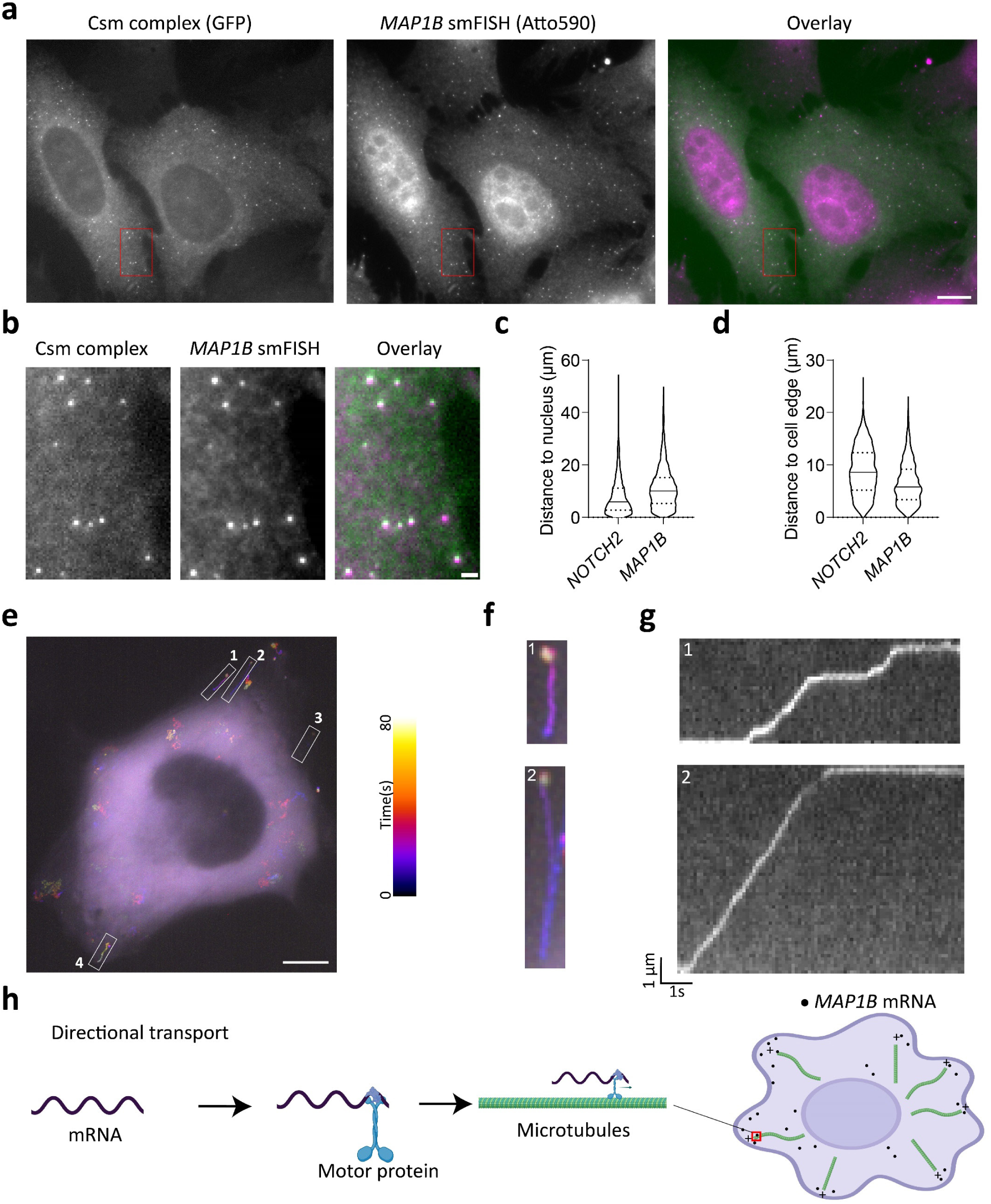
Dynamics of individual *MAP1B* mRNAs in live cells. **a**, Fixed-cell image of individual *MAP1B* mRNAs labeled by GFP-tagged Csm complex and 48 *MAP1B*-targeting spacers (left), image of individual *MAP1B* mRNAs labeled by smFISH probes (middle), and their overlay (right). Scale bar, 10 μm. **b**, Enlarged view of the red boxed region in **a**. Scale bar, 1 μm. **c,** Violin plot showing distance to the nucleus from individual *NOTCH2* or *MAP1B* mRNA molecules. Median indicated by solid line; quartiles indicated by dashed lines. **d,** Same as **c,** but showing distance to cell edge. **e**, Temporal-color coded trajectory of *MAP1B* mRNA in live U2OS cell. Scale bar, 10 μm. White boxed regions indicate 4 directional transport *MAP1B* mRNA molecules **f**, Enlarged views of the white boxed regions 1 and 2 in **e**. Full movie shown in Supplementary Movie 2. **g**. Kymograph of boxed regions 1 and 2 in **e** showing the directed movement of the indicated *MAP1B* mRNAs. **h**, A proposed directional transport mechanism directing *MAP1B* mRNA to cell periphery based on motor protein trafficking along microtubules.

To elucidate the mechanism that enriches *MAP1B* mRNAs at the cell periphery, we performed live-cell imaging. Strikingly, unlike *NOTCH2*, we frequently observed linear transport of *MAP1B* mRNAs toward the cell edge. Upon reaching the cell periphery, they tended to remain static (Fig. 3 e-g, Supplementary Movie 2). Occasionally, *MAP1B* mRNAs moved backward toward the cell nucleus but then eventually progressed again to the cell edge (e.g. region of interest [ROI] 2 in Fig. 3 e-g, Supplementary Movie 2). By analyzing these movement trajectories through kymograph, we also observed pausing of *MAP1B* mRNAs during their directional transport (Fig. 3g, Supplementary Movie 2). Previous mRNA quantification results from immunoprecipitation of kinesin 1 demonstrate interactions between microtubule motor-protein kinesin-1 and *MAP1B* mRNAs^36^. Taken together, these results suggest that, unlike perinuclear *NOTCH2* mRNA, *MAP1B* mRNA utilizes directional transport on microtubules as a driving force to localize to the cell edge (Fig. 3h).

### Translation inhibition induces P-body formation without affecting *MAP1B* mRNA transport

Most transported mRNAs are thought to be translationally inactive until reaching their destination for local translation^37^, but a recent study showed that translation can also occur before or during transport^38^. Especially given that *MAP1B* mRNA encodes a microtubule-associated protein that may explain its transport along the cytoskeleton, we asked whether inhibiting translation influences its transit and/or post-destination dynamics.

To test this, we treated cells with puromycin and performed smLiveFISH for *MAP1B* mRNA. The presence of puromycin did not change the observed directional transport of *MAP1B* mRNA towards the cell edge (Fig. 4a-c, Supplementary Movie 3), suggesting that movement is not coupled to translation. In fact, the mean transit speed of *MAP1B* mRNAs increased slightly but significantly from 1.3 μm/s to 1.5 μm/s following puromycin treatment (Fig. 4d). This might result from loss of polysomes and/or RNA-binding proteins from *MAP1B* transcripts, a possibility that remains to be tested. Similarly, we observed no change in dynamics of the stationary *MAP1B* mRNA population already at the cell edge upon puromycin treatment--compared to the obvious shift from slow to fast movement seen for *NOTCH2* mRNA (Fig. 4e, Supplementary Movie 3). Thus, unlike *NOTCH2* mRNA, *MAP1B* mRNA localization dynamics are not translation-dependent.

**Fig. 4:**
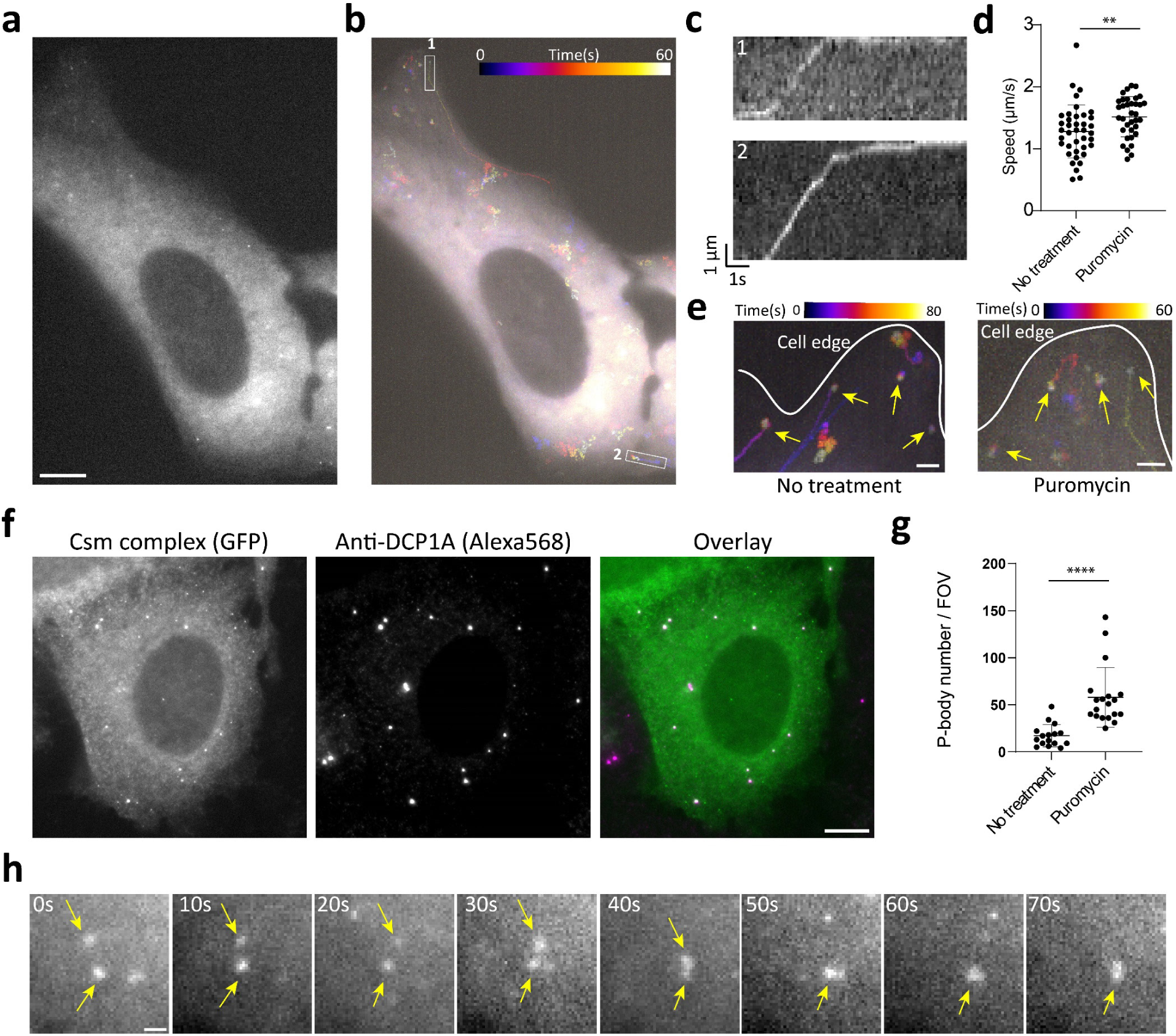
Dynamics of individual *MAP1B* mRNAs after puromycin treatment in live cells. **a**, Single snapshot of individual *MAP1B* mRNAs in live U2OS cell treated with puromycin. Scale bar, 10 μm. **b**, Temporal-color coded trajectory of *MAP1B* mRNA in cell shown in **a**. **c**, Kymograph of boxed regions 1 and 2 in **b** showing the directed movement of the indicated *MAP1B* mRNAs. Full movie shown in Supplementary Movie 3. **d**, Directed movement speeds of *MAP1B* mRNAs under untreated (n=39 events) and puromycin-treated (n=35 events) conditions. Each dot represents a directed movement event. **p < 0.01, t-test. **e**, Temporal-color coded trajectory of *MAP1B* mRNA at cell edge. Yellow arrows mark stationary *MAP1B* RNAs at the cell edge. Scale bar, 2 μm. **f**, Fixed-cell image of *MAP1B* mRNAs labeled by GFP-tagged Csm complex after puromycin treatment (left), immunostaining for the P-body marker DCP1A (middle), and their overlay (right). Scale bar, 10 μm. **g**, Quantification of P-body number with or without puromycin treatment. In both conditions, cells are seeded at the same density; each dot represents P-body number per field of view (FOV) with the same area. ****p < 0.0001, t-test. **h**, Time-lapse micrographs of *MAP1B* RNA granule formation after puromycin treatment in live U2OS cell. Yellow arrows mark two small puncta fusing into one larger RNA granule. Full movie is shown in Supplementary Movie 4. Scale bar, 1μm.

Interestingly, some *MAP1B* mRNAs coalesced into larger granules following puromycin treatment (Fig. 4f, Supplementary Movie 3). Because puromycin treatment has been shown to induce P-body formation and enlargement in U2OS cells^39^, we tested whether these RNA granules represent P-body formation. Co-staining for the P-body marker DCP1A showed that P-body number increased after puromycin treatment and that *MAP1B* mRNA granules colocalize with P-bodies (Fig. 4f-g). We noted one example of two small RNA puncta randomly moving around, contacting one another and eventually fusing into a single large granule (Fig. 4h, Supplementary Movie 4). Together these data show that smLiveFISH can be used to study RNA storage and metabolism in living cells.

## Discussion

SmLiveFISH provides a robust and programmable platform to directly visualize individual endogenous mRNAs in live cells using two components, a Csm protein-encoding plasmid and a programmable crRNA transcript-encoding plasmid. This platform enables real-time tracking of unmodified endogenous mRNAs at single-molecule resolution, revealing both spatial and temporal transcript dynamics in living cells. We developed a generalizable method to construct long crRNA arrays that produce the precursors for up to 24 guide RNAs complementary to a target RNA. Using smLiveFISH, we identified co-translational translocation and directional transport as two distinct methods for *NOTCH2* and *MAP1B* mRNA localization to the perinuclear and peripheral regions of the cell, respectively.

In parallel with smLiveFISH, we tested two Cas13 systems (PspCas13b and RfxCas13d) that have demonstrated imaging utility for highly abundant, repetitive RNAs^14, 40^. Both Cas13 proteins had limited efficacy in our hands even when attempting to image the abundant and repetitive XIST lncRNA (Extended data Fig. 1). The comparative effectiveness of CRISPR-Csm for single-transcript imaging shown in this study may result from Csm’s ∼30-fold higher binding affinity for RNA compared to Cas13^22,26^, and/or its multi-subunit nature, allowing for ≥3 Csm3-GFP molecules per complex. Experiments using MS2-tagged mRNA reporters have suggested that mRNA associations with endosomes, lysosomes and mitochondria account for localized delivery, translation and co-translational protein complex assembly^30, 41^. SmLiveFISH will enable exploration of RNA-organelle interactions by allowing direct observation of unmodified endogenous mRNAs.

Many neurodegenerative disorders are caused by mutations in RNA-binding proteins. Popular examples include mutations in FMRP responsible for Fragile X Syndrome (FXS) and in TDP-43 responsible for Amyotrophic lateral sclerosis (ALS). Both FMRP and TDP-43 have been shown to regulate *MAP1B* mRNA transport, translation, and/or stability in neurons^42–44^, and dysregulation of *MAP1B* and other target mRNAs has been proposed as a contributing factor in the disease^45^. SmLiveFISH can now be used as a sensitive assay to explore such mechanisms in ways that fixed-cell assays cannot, such as by measuring changes in transport speed or step-wise displacement (as done in Figs. 3-4). This may help uncover the pathological mechanisms of RNA-centric diseases such as FXS and ALS.

We observed different behaviors of *NOTCH2* and *MAP1B* mRNA after puromycin treatment, with *MAP1B* but not *NOTCH2* mRNA forming large RNA granules coincident with P-bodies. Previous sequencing results for mRNAs isolated from purified P-bodies support these findings^46^, whereby mRNAs encoding integral ER proteins are depleted in P-bodies, while those encoding proteins involved in cell division, differentiation and morphogenesis are enriched^46^.

These findings suggest that *NOTCH2* and *MAP1B* mRNAs may rely on different decay pathways whose sorting mechanisms have yet to be elucidated.

Structural investigation of type III CRISPR-Cas systems indicates that the Csm1 and Csm4 subunits recognize the (−6) and (−7) nucleotides of the crRNA 5′ handle in a sequence-specific way^25^. Thus, it may be possible to develop orthogonal type III CRISPR-Cas systems with minimal crosstalk to the type III-A CRISPR-Csm complex from *S*. *thermophilus* used here, enabling multi-color live-cell RNA imaging. This method can then be extended to address questions about RNA-RNA interactions, RNA splicing and co-translational protein complex assembly. We expect smLiveFISH to accelerate efforts to study the spatiotemporal dynamics of various RNA species in many contexts, with immediate applications in the RNA, cell biology and neurobiology fields.

## Methods

### Cell lines and cell culture

HEK293T, U2OS, HeLa, IMR-90 and COS-7 cells were obtained from the UC Berkeley Cell Culture Facility, and were grown in medium containing DMEM, high glucose (Thermo Fisher Scientific), 10% FBS (Sigma), and 1× Pen/Strep (Thermo Fisher Scientific) at 37°C with 5% CO2.

### Plasmid construction

Nuclear-targeting Csm complex plasmid construction has been described previously^12^ (Addgene plasmid #195242). Cytoplasmic-targeting Csm complex plasmid was modified from this by removing the NLS sequences before each protein sequence, changing dCsm-EGFP to dCsm-2xsfGFP, and removing the U6-crRNA region. For the crRNA array plasmid, *NOTCH2* arrays were constructed through multiple steps of overlap extension PCR, as described in Extended Data Fig. 2. *MAP1B* arrays were constructed similar to *NOTCH2*, but the overlapping regions were implemented in the oligo pool sequences to skip the step3 (Supplementary Table 1). Arrays were cloned after a CAG promoter with a short signal sequence^27^ from the HSPB3 gene placed between the CAG promoter and crRNA array to help export from the nucleus to the cytoplasm. All cloning was performed in NEB stable E. coli (NEB) to prevent recombination between repetitive sequences. Plasmids were verified by whole-plasmid sequencing. dPspCas13b-3xEGFP plasmid was purchased from Addgene (plasmid #132398), dRfxCas13d-EGFP was modified from Addgene plasmid #109050 by removing the T2A sequence between dRfxCas13d and EGFP. CrRNA and oligo pool sequences are listed in Supplementary Table 1. Plasmid sequences are listed in Supplementary Table 2.

### Optical setup and image processing

Cell samples were imaged using a wide-field fluorescent microscope (Zeiss Axio Observer Z1 inverted fluorescence microscope) with a 100×/1.4 NA oil Ph3 Plan Apochromat objective, an ORCA-Flash4.0 camera (Hamamatsu), a X-Cite 120Q lamp, and ZEN 2012 software. GFP filter sets include the BP 470/40 excitation filter, the FT 495 beamsplitter, and the BP 525/50 emission filter. Atto 590 and Alexa Fluor 568 filter sets include the BP 572/25 excitation filter, the FT 590 beamsplitter, and the BP 629/62 emission filter. Images representing max-intensity z-projections were generated by FIJI software. Colocalization analysis was performed by FIJI plugin, ComDet V.0.5.5. Single-molecule tracking was performed by FIJI plugin, TrackMate. The temporal-color coded images (Fig. 3e-f, 4b, 4e) were generated using FIJI Temporal-Color Code function. To generate kymographs in Fig. 3g and 4c, polyline selections were used to track the particle moving trajectories, then kymographs were generated using KymoResliceWide plugin.

### Immunostaining

For H2AK119ub staining, 150K HEK293T cells were grown on 18mm diameter, #1.5 thickness, collagen coated coverslips (Neuvitro) in a 12-well plate. The next day, cells were transfected with 0.8 μg *XIST-*targeting dCsm-EGFP complex plasmid DNA, 0.4 μg dPspCas13b-3xEGFP plus 0.6 μg *XIST*-targeting PspCas13b crRNA plasmid, or 0.4 μg dRfxCas13d-EGFP plus 0.6 μg *XIST-*targeting RfxCas13d crRNA plasmid using 5 μl TransIT-293 transfection reagent (Mirus Bio). After transfection, cells were grown for 48 hours to allow plasmid expression. Then, cells were fixed with 4% paraformaldehyde (Electron Microscopy Sciences) in 1× PBS at room temperature for 10-15 min. Following three washes with 1× PBS, cells were permeabilized by 0.5% (vol/vol) Triton X-100 (Sigma) in 1× PBS for 10 min at room temperature. Samples were again washed with 1× PBS three times after permeabilization. The permeabilized cells were incubated in blocking buffer (1xPBS containing 3% (wt/vol) BSA (Jackson ImmunoResearch)) for 1 hour. Cells were then incubated with H2AK119ub primary antibodies at 1:1000 dilution (Cell Signaling) in blocking buffer for 1 h at room temperature and washed with 1× PBS three times for 5 min each. Next, cells were stained with Alexa Fluor 568-labeled secondary antibodies in blocking buffer for 1 h at room temperature. Samples were washed again with 1× PBS three times to remove unbound antibodies. To prevent bound antibody dissociation, samples were post-fixed with 4% (vol/vol) PFA in 1× PBS for 10 min and washed three times with 1× PBS for 5 min each.

For DCP1A staining, 100k U2OS cells were grown on 18mm diameter, #1.5 thickness, collagen coated coverslips (Neuvitro) in a 12-well plate. The next day, 0.8 μg cytoplasmic-targeting Csm complex plasmid and two *MAP1B* crRNA array plasmids (0.7 μg each) were transfected into cells using 5 μl TransIT-LT1 transfection reagent (Mirus Bio). After transfection, cells were cultured for 48 hours to allow plasmid expression. Antibody staining was performed as for the above H2AK119ub procedure, but with anti-DCP1A antibody (Abcam).

## RNA FISH

HEK293T, HeLa, IMR-90 and COS-7 cells were grown on 18mm diameter, #1.5 thickness, collagen coated coverslips (Neuvitro) in a 12-well plate. Next day, 0.8 μg cytoplasmic-targeting Csm complex plasmid and two crRNA array plasmids (0.7 μg each) were transfected into cells using 5 μl TransIT-293 transfection reagent (Mirus Bio) or TransIT-LT1 transfection reagent (Mirus Bio). For U2OS cells, 1x10^6^ cells were nucleofection with 1.5 μg cytoplasmic-targeting Csm complex plasmid and two crRNA array plasmids (1.2 μg each). Then U2OS cells were seeded on 18mm diameter, #1.5 thickness, collagen coated coverslips (Neuvitro) at 250k density per well. After transfection, cells were grown for 48 hours to allow plasmid expression. Then, cells were fixed with 4% paraformaldehyde (Electron Microscopy Sciences), followed by permeabilization with 0.5% (vol/vol) Triton X-100 (Sigma) in 1× PBS for 10 min at room temperature. Next, cells were incubated for 5 min in wash buffer comprising 2×SSC (Thermo Fisher Scientific), 30% (vol/vol) formamide (Thermo Fisher Scientific), and then stained with *NOTCH2* or *MAP1B* mRNA FISH probes in hybridization buffer containing 30% (vol/vol) formamide, 0.1% (wt/vol) yeast tRNA (Thermo Fisher Scientific), 1% (vol/vol) murine RNase inhibitor (New England Biolabs), 10% (wt/vol) dextran sulfate (Sigma), and 2×SSC in a humidity-controlled 37 °C incubator overnight. FISH probes were stained at a concentration of 200 nM (5nM per probe, ∼40 probes in total). After staining, cells were washed twice with wash buffer at 37 °C, each for 30 minutes. Then, cells were stained with DAPI and 5 nM readout probes in separate hybridization buffer composed of 2× SSC, 10% (vol/vol) ethylene carbonate (Sigma) in nuclease-free water before imaging. The *NOTCH2* and *MAP1B* mRNA FISH probes sequences and readout probe sequence are provided in Supplementary Table 3. *NOTCH2* and *MAP1B* FISH probes are ordered as oligo pools from IDT.

### Live Cell Imaging

For live cell imaging of *NOTCH2* and *MAP1B* mRNA, 1x10^6^ U2OS cells were nucleofected with 1.5 μg cytoplasmic-targeting Csm complex plasmid and two crRNA array plasmids (1.2 μg each). Then, U2OS cells were seeded in a 2-well glass bottomed NuncLab-Tek chamber (Thermo Fisher Scientific) at 400k density per well. After 48 hours, the medium was changed to live cell imaging buffer containing no-phenol red medium supplied with 10% FBS, 1× Pen/Strep, and ProLong live antifade reagent (Thermo Fisher Scientific).

### Puromycin treatment

For puromycin treatment, cells were incubated in live-cell imaging buffer containing 275 μM puromycin for 60 minutes at 37°C prior to fixation or live-cell imaging.

### SMdM Data analysis

SMdM analyses are described previously^34^. Briefly, single-molecule spots were first localized in all frames. Paired locations were identified across successive frames for calculation of displacements in the frame time Δt = 100 ms. The displacements were spatially binned with a grid size of 2.5 pixels (325 nm). The displacements in each spatial bin were separately fitted to a single-component diffusion mode through maximum likelihood estimation:

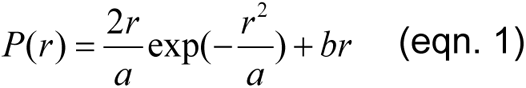

Here *a* = 4*D*Δ*t* with *D* being the diffusion coefficient, and *b* accounts for a uniform background. The resultant local apparent *D* values were presented on a continuous color scale to produce a diffusivity map (Fig. 2c, d). Separately (Fig. 2e, f), all single-molecule displacements in each cell were pooled and fitted to a two-component diffusion mode^47^:

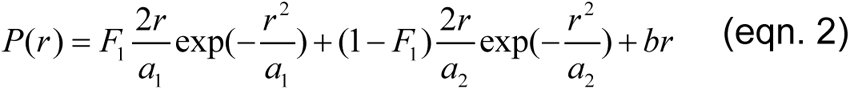

where *F*_1_ and *F*_2_ = (1 − *F*_1_) are the fractions of the two diffusivity components, and *a*_1_ = 4*D*_1_Δ*t* and *a*_2_ = 4*D*_2_Δ*t* account for the two diffusion coefficients *D*_1_ and *D*_2_.

## Author contributions

C.X. and J.A.D. conceived of the project. J.A.D. supervised the project. D.C. conducted initial cytoplasmic knockdown tests using the Csm system. C.X. reconstructed and repurposed the Csm system for cytoplasmic RNA imaging. C.X. and X.J. developed the crRNA array construction method. C.X., X.J. designed and performed all the imaging experiments. C.X. and K.X. performed data analyses. C.X., K.X., and J.A.D. wrote the paper with input from D.C., X.J.

## Acknowledgements

We thank all members of the Doudna lab for helpful advice and discussions. This project was supported by HHMI and grants to J.A.D. from the NIH. D.C. was supported in part by the Jane Coffin Childs Memorial Fund for Medical Research. J.A.D. is an investigator of the Howard Hughes Medical Institute. K.X. acknowledges support by the National Institute of General Medical Sciences of the National Institutes of Health (R35GM149349).

## Competing interests

J.A.D. is a cofounder of Azalea Therapeutics, Caribou Biosciences, Editas Medicine, Evercrisp, Scribe Therapeutics, Intellia Therapeutics, and Mammoth Biosciences. J.A.D. is a scientific advisory board member at Evercrisp, Caribou Biosciences, Intellia Therapeutics, Scribe Therapeutics, Mammoth Biosciences, The Column Group and Inari. J.A.D. is Chief Science Advisor to Sixth Street, a Director at Johnson & Johnson, Altos and Tempus, and has a research project sponsored by Apple Tree Partners. C.X. and J.A.D. are inventors on patents applied for by the Regents of the University of California related to Csm complex single-molecule imaging.

---

**Extended Data Fig. 1:**
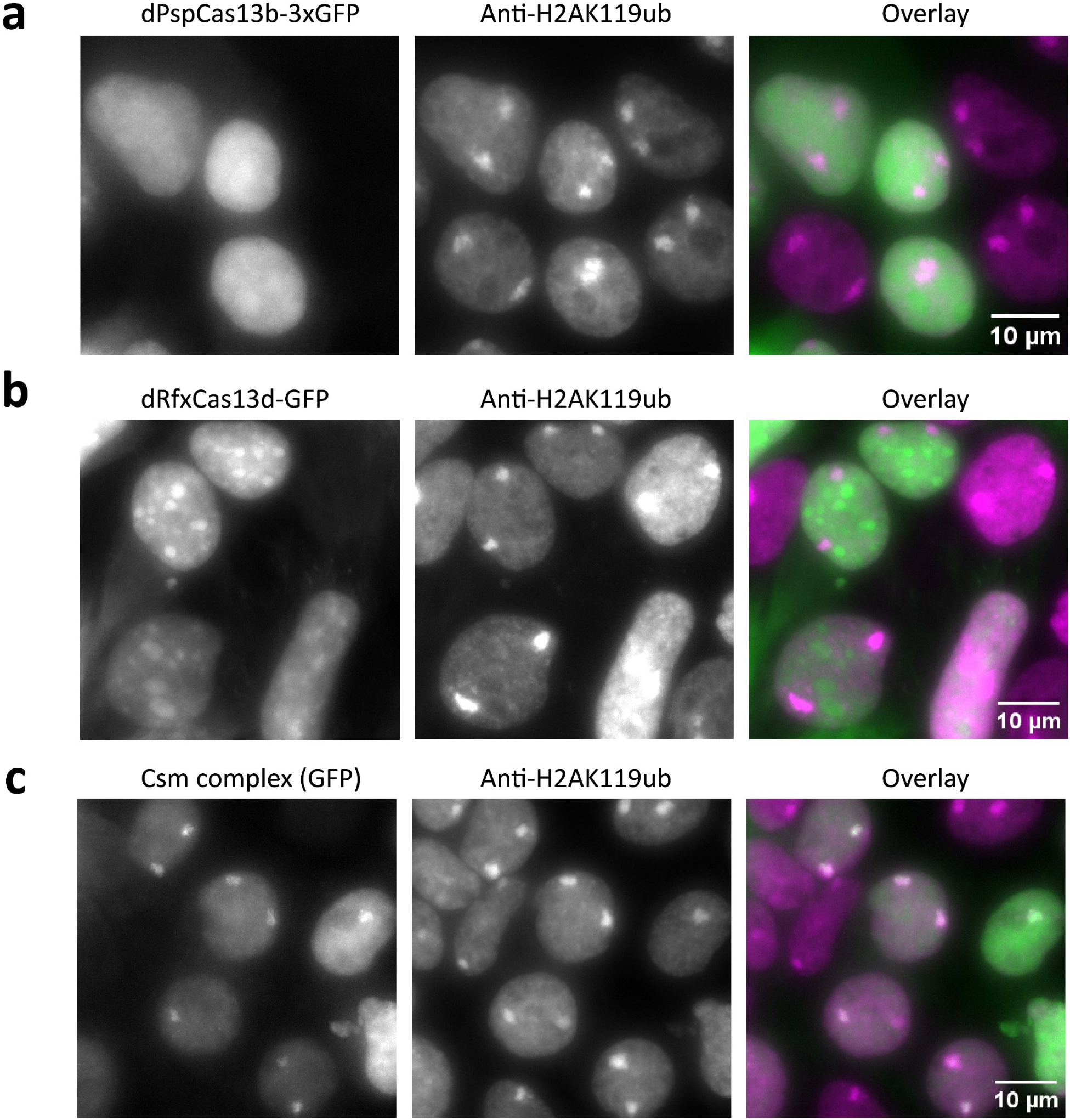
Testing different Cas proteins for robust RNA imaging. Representative images of GFP-tagged PspCas13b (**a**), RfxCas13d (**b**) and Csm complex (**c**) targeting *XIST* RNA in HEK293T cells (left), immunostaining for the heterochromatin marker H2AK119ub (middle), and their overlay (right). Scale bar, 10 μm.

**Extended Data Fig. 2:**
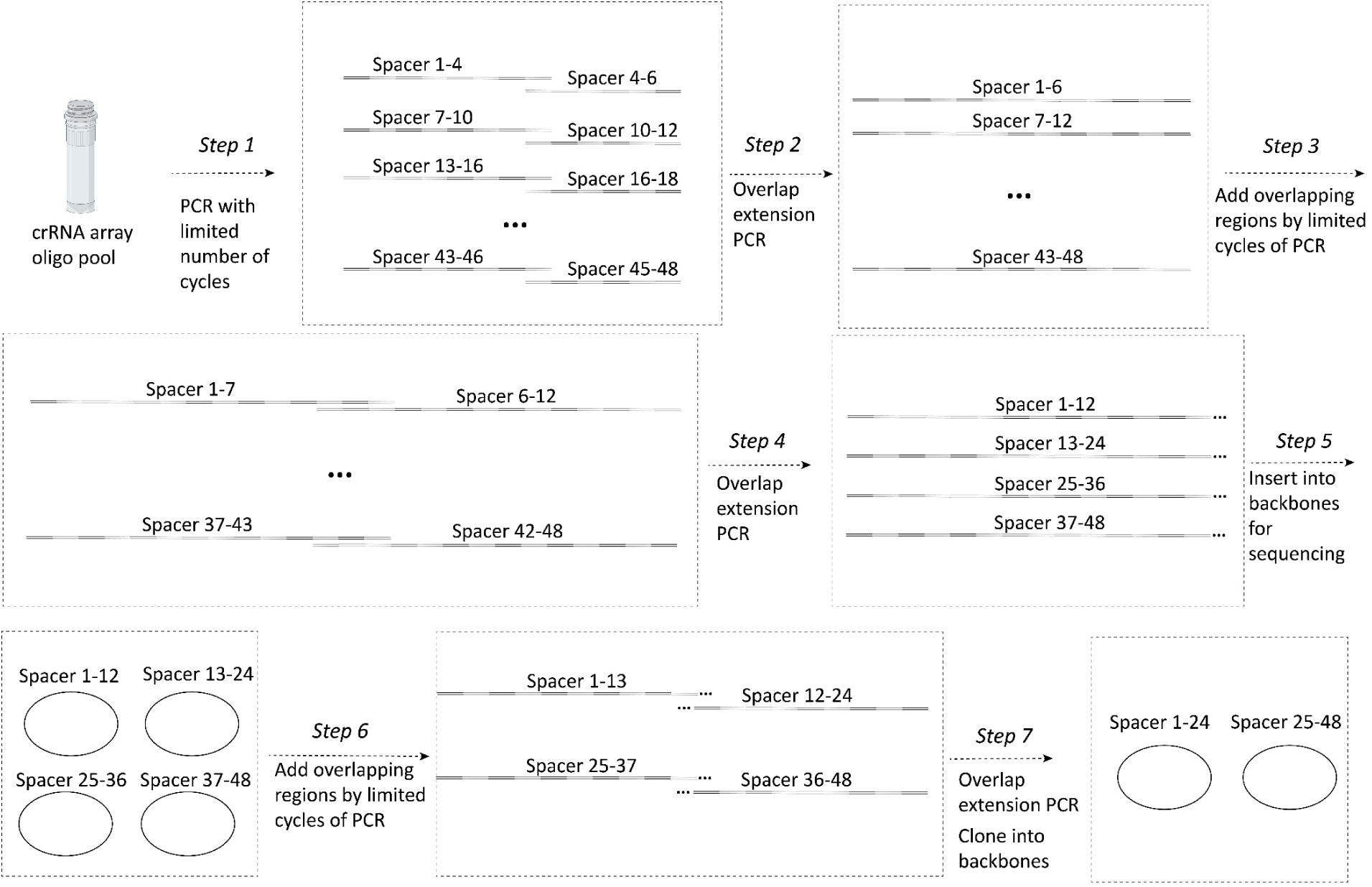
CRISPR array cloning method. Schematic of CRISPR array construction. First, each fragment from an oligo pool containing multiple 3-4 spacer fragments, is amplified by limited-cycle PCR. Second, 6-spacerfragments are generated by overlap extension PCR of the 3-4-spacer fragments. Third, overlapping ends are added to the 6-spacer fragments by limited-cycle PCR. Fourth, 12-spacer fragments are generated by overlap extension PCR of the 6-spacer fragments. Fifth, the 12-spacer fragments are cloned into backbones for sequence verification. Sixth, overlapping ends are added to the 12-spacer fragments by limited cycle PCR. Seventh, 24-spacer fragments are generated by overlap extension PCR of the 12-spacer fragments by overlap extension PCR. Finally, the 24-spacer fragments are cloned into backbones for sequence verification.

**Extended Data Fig. 3:**
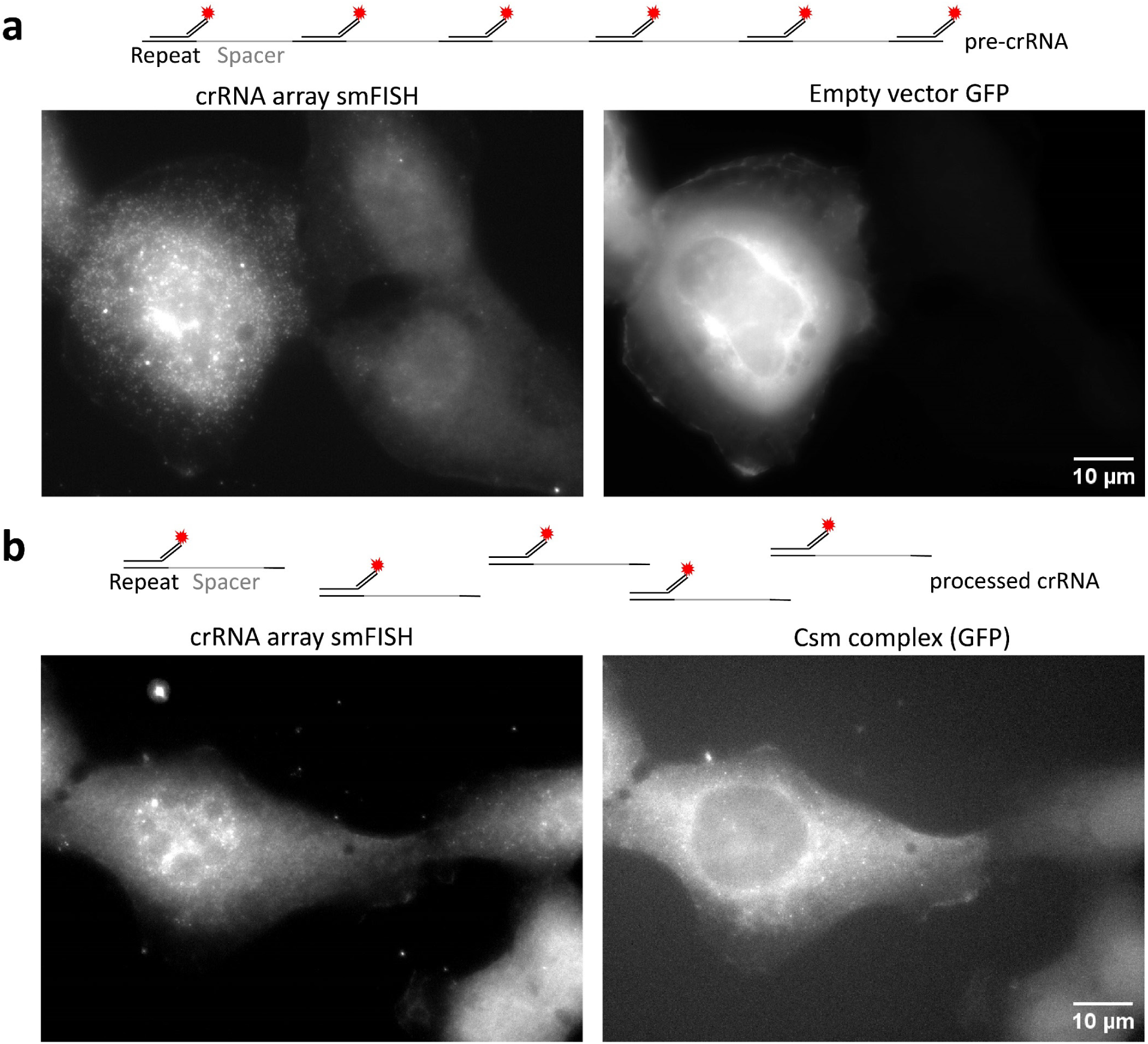
Verifying pre-cRNA processing in cells. **a**, Representative image of smFISH against crRNA array transcripts in U2OS cells co-transfect with plasmid expressing CRISPR array and GFP alone. Left, crRNA array smFISH spots detected by a single FISH probe (Atto590) that binds the direct repeats of the crRNA array. Right, GFP channel for the same cells in the left panel. Scale bar, 10 μm. **b**, Representative image of smFISH against crRNA array transcripts in U2OS cells co-transfected with plasmid expressing CRISPR array and GFP-tagged Csm complex. Left, crRNA array smFISH spots detected by a single FISH probe (Atto590) that binds the direct repeats of the crRNA array. Right, GFP channel for the same cells in the left panel. Scale bar, 10 μm.

**Extended Data Fig. 4:**
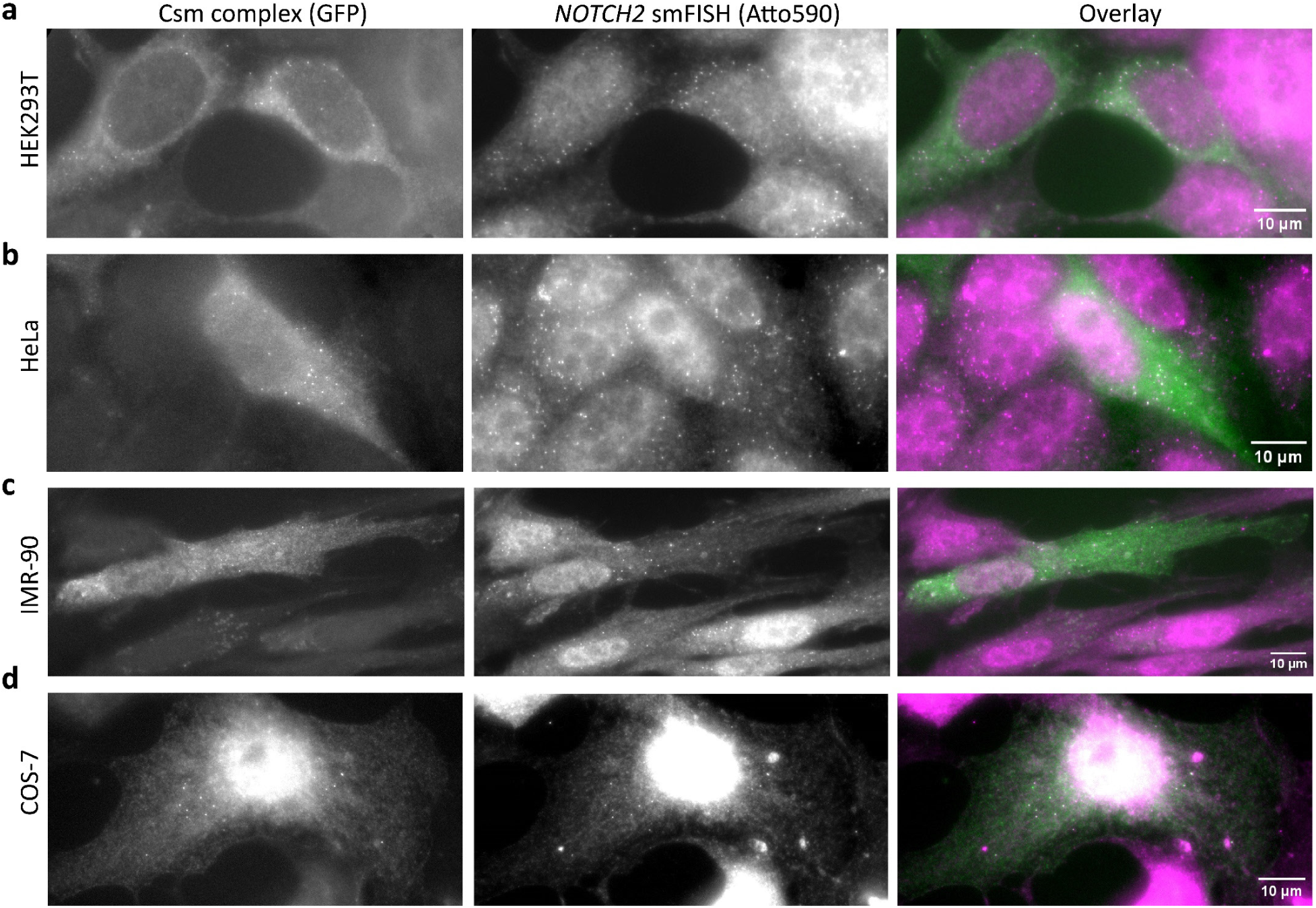
smLiveFISH works across various cell types. SmLiveFISH image of individual *NOTCH2* mRNAs labeled by GFP-tagged Csm complex (left), smFISH image of individual *NOTCH2* mRNAs (Atto590), and their overlay (right) in HEK293T (**a**), HeLa (**b**), IMR-90 (**c**), and COS-7 cells (**d**). Scale bar, 10 μm.

**Extended Data Fig. 5:**
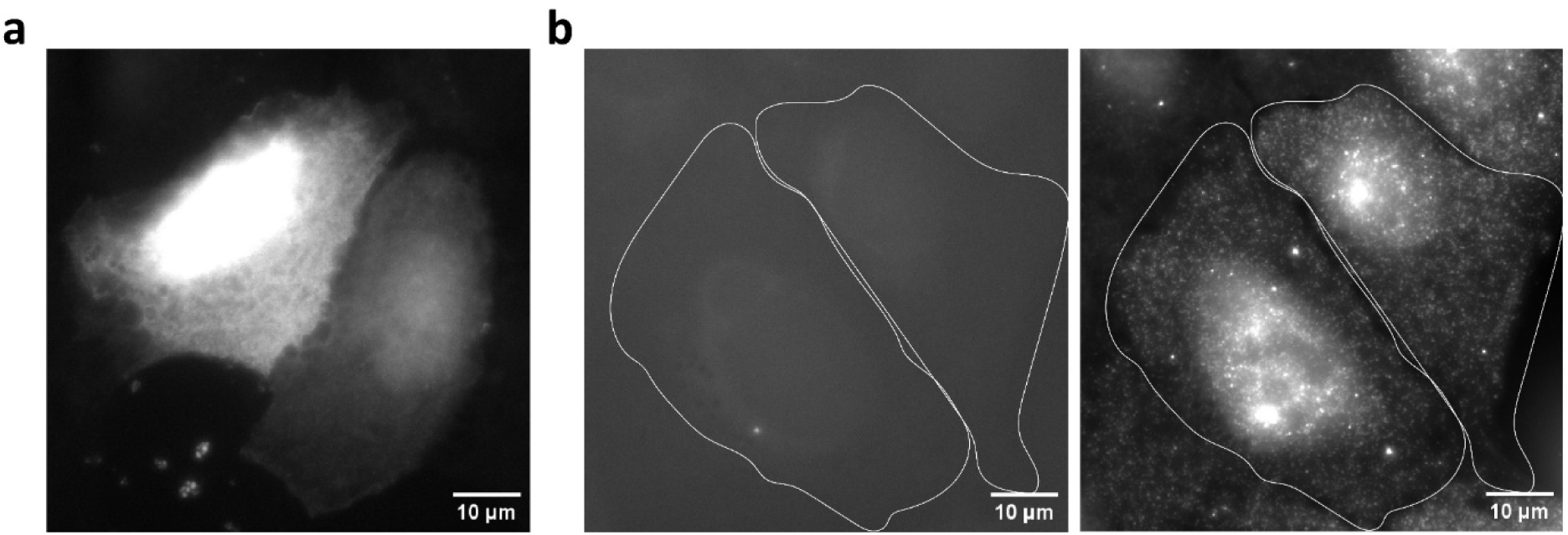
Imaging control for RNA labeling with Csm complex. Representative GFP channel signal of cells expressing GFP-tagged Csm complex plasmid alone (**a**) or crRNA array plasmid alone (**b,** left). Right panel of **b** shows smFISH image of the crRNA array labeled by Atto590. Scale bar, 10 μm.

**Supplementary Movie 1**: *NOTCH2* mRNA visualized 60 min after addition of puromycin. Compared to no treatment (left), movement of *NOTCH2* mRNA shifts from the slow to the fast movement category. Movie recorded at 10 frames per second. Scale bar, 5 μm.

**Supplementary Movie 2**: Live-cell imaging of *MAP1B* mRNA in untreated U2OS cells. Movie recorded at 10 frames per second. Scale bar, 10 μm.

**Supplementary Movie 3**: Live-cell imaging of *MAP1B* mRNA in puromycin-treated U2OS cells. Movie recorded at 10 frames per second. Scale bar, 10 μm.

**Supplementary Movie 4**: Live-cell imaging of *MAP1B* mRNA in puromycin-treated U2OS cells. The white boxed region shows *MAP1B* RNA granule formation. Movie recorded at 10 frames per second. Scale bar, 10 μm.

